# Measuring popularity of ecological topics in a temporal dynamical knowledge network

**DOI:** 10.1101/474148

**Authors:** Tian-Yuan Huang, Bin Zhao

## Abstract

As interdisciplinary branches of ecology are developing rapidly in the 21^st^ century, contents of ecological researches have become more abundant than ever before. Along with the exponential growth of number of published literature, it is more and more difficult for ecologists to get a clear picture of their discipline. Nevertheless, the era of big data has brought us massive information of well documented historical literature and various techniques of data processing, which greatly facilitates the implementation of bibliometric analysis on ecology. Frequency has long been used as the primary metric in keyword analysis to detect ecological hotspots, however, this method could be somewhat biased. In our study, we have suggested a method called PAFit to measure keyword popularity, which considered ecology-related topics in a large temporal dynamical knowledge network, and found out the popularity of ecological topics follows the “rich get richer” and “fit get richer” mechanism. Feasibility of network analysis and its superiority over simply using frequency had been explored and justified, and PAFit was testified by its outstanding performance of prediction on the growth of frequency and degree. In addition, our research also encourages ecologists to consider their domain knowledge in a large dynamical network, and be ready to participate in interdisciplinary collaborations when necessary.

## 1. Introduction

Early in 1994, historian Donald Worster had made an interesting remark in his book, “Ecology achieved intellectual sophistication, academic prominence, and financial security in the postwar years, but also lost much of its coherence. It broke down into a cacophony of subfields, including ecosystematists, populationists, biospherians, theoretical modelers, forest and range managers, agroecologists, toxicologists, limnologists, and biogeographers”(Worster 1994). By now, this remark still stands and could not be more correct. The scope of ecological research is expanding unprecedentedly in 21^st^ century. Relations between biological systems and surrounding environments are of great complexity, numerous disciplines are joining ecology to answer demanding ecological questions and meet the global challenge. This has opened a door for discipline integration, and various branches of ecology had emerged in recent decades, with new theories, methods and technologies (Thompson *et al.* 2001). As the number of ecological literature is growing faster and faster in recent years(Nunez Mir *et al.* 2016), it is becoming more and more difficult for ecologists to get a clear picture of knowledge structure in their study area, not to mention the broad overview of the whole discipline.

But thanks to the era of big data, it is now getting easier and easier for scientists to get mass literature data. Together with the handy tools from automated content analysis, scientists can now carry out bibliometric research and dig deep into the historical ecological literature. (Nunez Mir *et al.* 2016; Kim *et al.* 2018). In this way, new insights on the trends of ecology could be discovered in novel ways. This could be an excellent complement to the traditional literature overview.

In bibliometric studies, keyword analysis, as core content summary of articles, has long been used to identify research focus in ecological disciplines (Budilova *et al.* 1997; Liu *et al.* 2011; Song & Zhao 2013; Stork & Astrin 2014; Wang *et al.* 2015; Romanelli *et al.* 2018). Author keywords contain information that authors consider as most concerned and relevant to their studies, and high-frequency keywords are deemed to reflect the hot issues, and could be used to reveal the research trends (Li *et al.* 2011; Li *et al.* 2017; Yang *et al.* 2017; Yin *et al.* 2018). Usually, keywords are ranked according to their frequency and sorted in a descending order, high ranking keywords are showed in a list, and we get an overview of the research hotspots from these most frequently used author keywords. By implementing the above method, it is already assumed that topics behind high-frequency keywords are more popular than others.

We have doubts about this assumption, for a topic is not only popular for frequently occurring in literatures, but also for it could be widely accepted in public and co-occurred with various other topics in the same article. Previous studies have applied co-word analysis to address this problem (Zhuang *et al.* 2013; Wang *et al.* 2015; Chen *et al.* 2016; Aleixandre-Benavent *et al.* 2018). Using keyword co-occurrence network, the relationships of keywords could be depicted, and the centrality of keywords could be vividly showed. Nevertheless, most co-word analyses were restricted to simple descriptions of the network, few studies dig deep into the application of social network analysis, and quantitative studies were seldom carried out to further explore the trends of ecology. Therefore, most of the times frequency is still the only metric to measure keyword popularity in bibliometric analysis.

To fill this gap, we first constructed the ecological knowledge network with 247,764 articles from 137 leading ecological journals based on the co-occurrence of author keywords. Then we asked research questions as follows: Is network analysis feasible to detect hotspots in ecology? What are the possible risks when using frequency to measure keyword popularity compared with network-based methods? When the previous questions were answered, we proposed an approach called PAFit, which had been applied successfully in the research of scientific collaboration (Ronda-Pupo & Pham 2018), to measure keyword popularity in a temporal dynamical network. In the proposed method, the keywords in ecological journals were considered as ecology-related topics, and tested to see if they follow “rich get richer” and “fit get richer” mechanism. At last, our proposed method was testified by a comparative study. The main objective of our work was to propose a new method to measure keyword popularity. But other than this, we hoped our study could encourage ecological researchers to consider their domain knowledge in a broad network, and be ready to join transdisciplinary researches while focusing on their specific studies.

## 2. Materials and Methods

### 2.1 Data source

To build a comprehensive database of ecological literature information, we consulted the latest ISI Journal Citation Reports (2017) and chose journals under the “ecology” category (more details could be found in S1 Table). The information of ecological journals was downloaded from SCOPUS (https://www.scopus.com), where we could export at most 2,000 documents per time in csv format efficiently. For the reason that digital archives of historical data were not so complete in the 1900s, we limited our time range to the recent 30 years, namely from 1988 to 2017. Also, only papers with document type of “article” were chosen, and entries containing missing values were excluded in our database. As keywords are not case-sensitive, all the keywords were converted to lower case, and duplicated records were merged. After data cleaning, we finally got a dataset with 247,764 papers from 137 leading ecological journals (detailed names of journals could be found in S1 Table). The annual article number was increasing steadily in our dataset, which led to the bursting number of distinct keywords that poured into the ecological disciplines (Fig.1). Since these articles came from journals categorized as “ecology”, keywords in these articles were considered to be relevant with ecology. Therefore, these keywords possess the potential to become ecological topics in the community of ecological researchers.

**Fig. 1.**
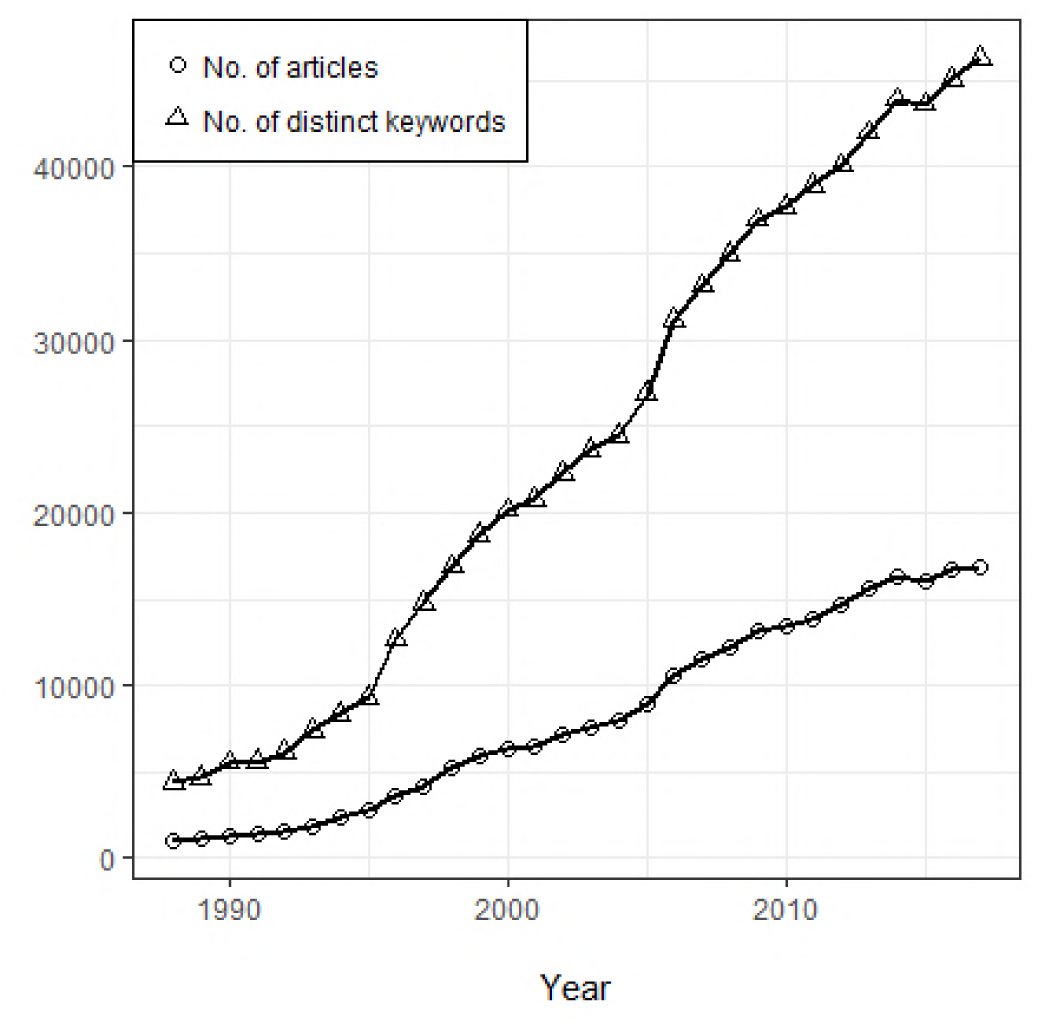
Annual article number and distinct keyword number based on our data source.

### 2.2. Construction of ecological knowledge network

To construct ecological knowledge network, we have a basic assumption that keywords co-occurred in the same article are related to each other. For a single article, when we get the keywords list, we could gain the keyword co-occurred relations among these keywords, which provide an edge list to construct the final network (Fig.2). We could find that keywords in the same article are all linked to each other in the network. When we had more papers, we could extract the keyword co-occurred relations from large amount of articles and formed a huge complex knowledge network (Fig.3). We believed this network could provide important information on knowledge structure of ecology and had the potential to detect and quantify ecological research hotspots. The whole network establishment procedure was conducted in R with packages including ‘igraph’(Csardi & Nepusz 2006), ‘ggraph’ (Pedersen 2017)and ‘tidygraph’(Pedersen 2018).

**Fig. 2.**
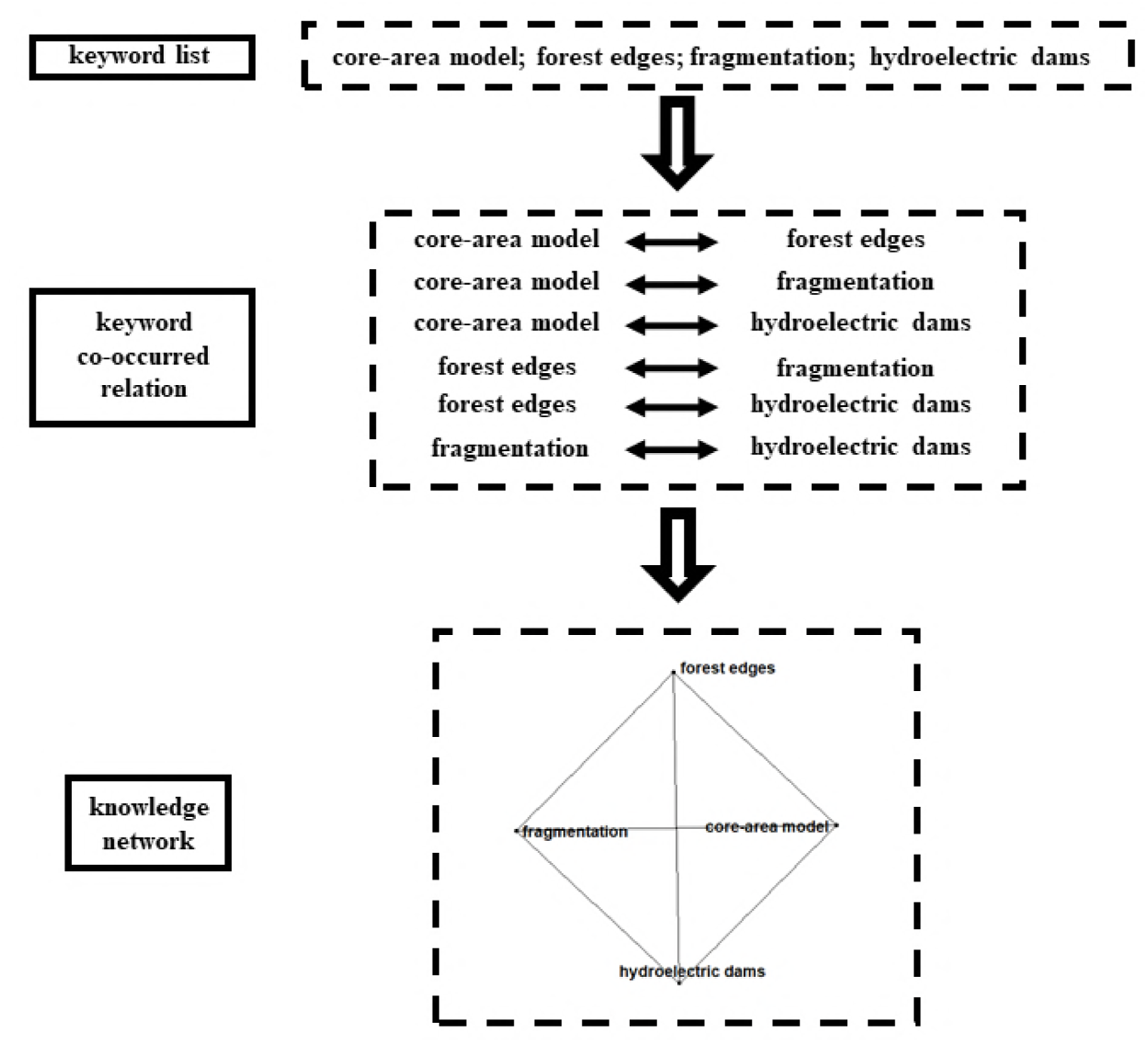
Construction of knowledge network from a single article. (The sample displayed here came from a real article published in *Acta Amazonica*. Ferreira *et al.* 2012)

**Fig. 3.**
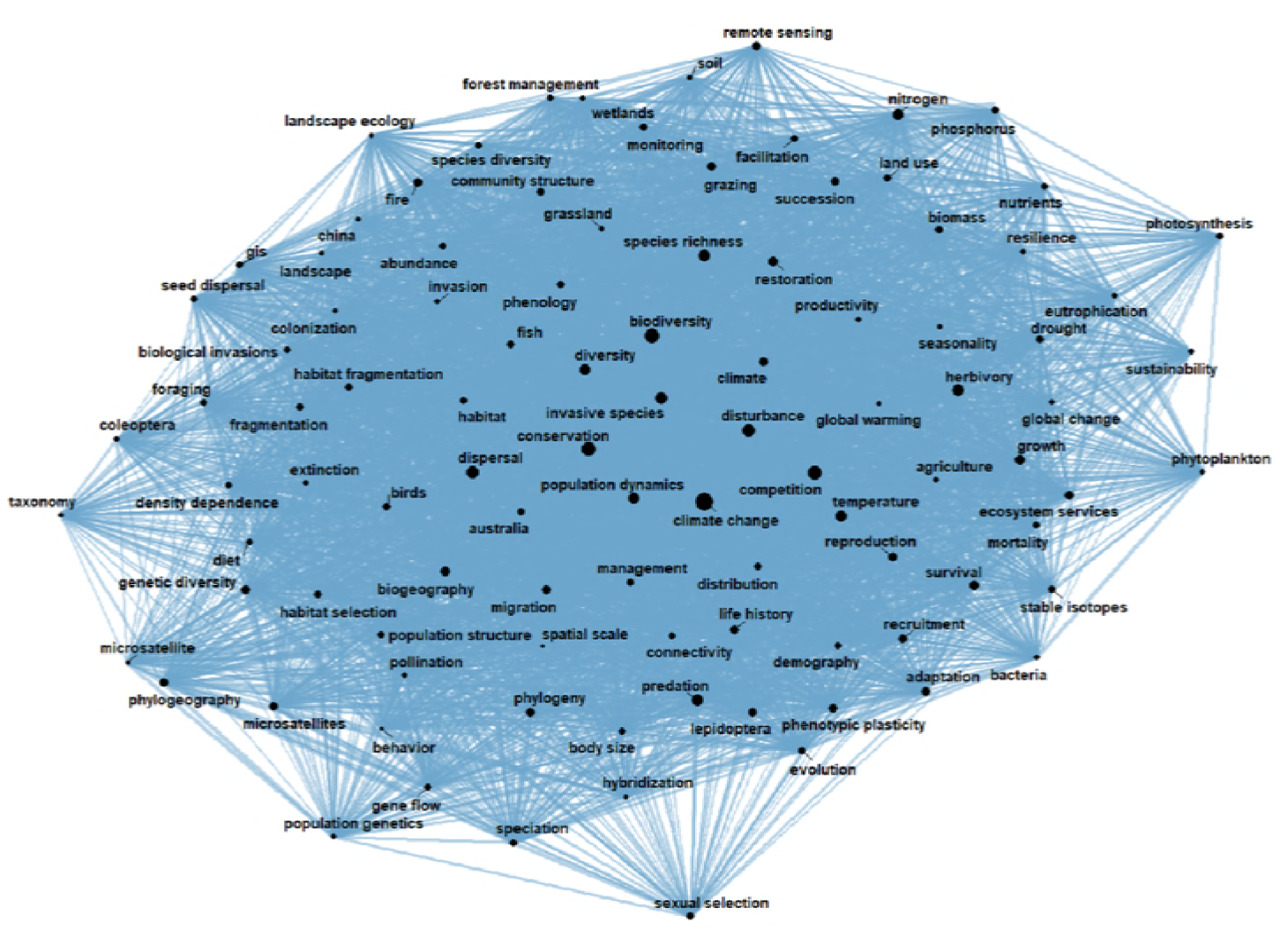
Ecological knowledge network. The above network is established from data covering 30 years (1988-2017), only 100 keywords with largest degree are displayed (the total network is an undirected graph with 312,767 nodes and 3,321,885 edges). The sizes of nodes are rescaled by the node degree, and the width of edges are proportional to the co-occurring times of the two keywords.

### 2.3. Interpretations of concepts from network analysis in our study

In graph theory, numerous metrics are used to describe network properties in different levels, including node-level, group-level and network-level (Al-Taie & Kadry 2017). Because we wanted to quantify the popularity of ecological topics, we had first chosen the simplest but maybe the most effective node-level centrality metric, degree. The degree of a node is the number of links it has with other nodes, therefore, the popularity of the node is determined by how many nodes it is connected to (Luke 2015; Al-Taie & Kadry 2017). When it comes to our study, degree of a keyword (represented by a node in the network) is the measure of the capability to co-occur with other keywords in the same article. As each keyword represents an ecology-related topic, the popularity of the topic could be reflected by how many different topics it could be related to.

We had also used network-level metric density to depict the compactness of the knowledge network. By definition, the density is the proportion of edges in the network to the maximum number of possible edges. As our network is undirected, the density **D (G) = 2m/ (n*(n-1))**, where **n** is the total node number and **m** is the total edge number.

### 2.4. Comparison of different results yielded by frequency and degree when measuring keyword popularity

We believed that degree calculated in the constructed knowledge network could be a good competitor against the commonly used metric frequency on the task of measuring keyword popularity, therefore we tried to find the difference in the results yielded by frequency and degree. First, we gathered all the keywords from ecological articles during the recent three decades, and calculated their frequency and degree. Then we ranked the keywords according to both metrics, which generated two different ranking lists. The differences between frequency ranking and degree ranking were calculated so we could find the main distinctions between them. Only top 1,000 keywords in degree ranking list or frequency ranking list were taken into consideration, so that keywords we selected had certain influences in ecology. At last we made two lists, one for keywords with relatively low frequency but high degree, the other for keywords with relatively high frequency but low degree. Geographical names like “france” and “oregon” were excluded and only 20 keywords with largest differences were shown in the lists (Table 1, Table 2).

**Table 1.**
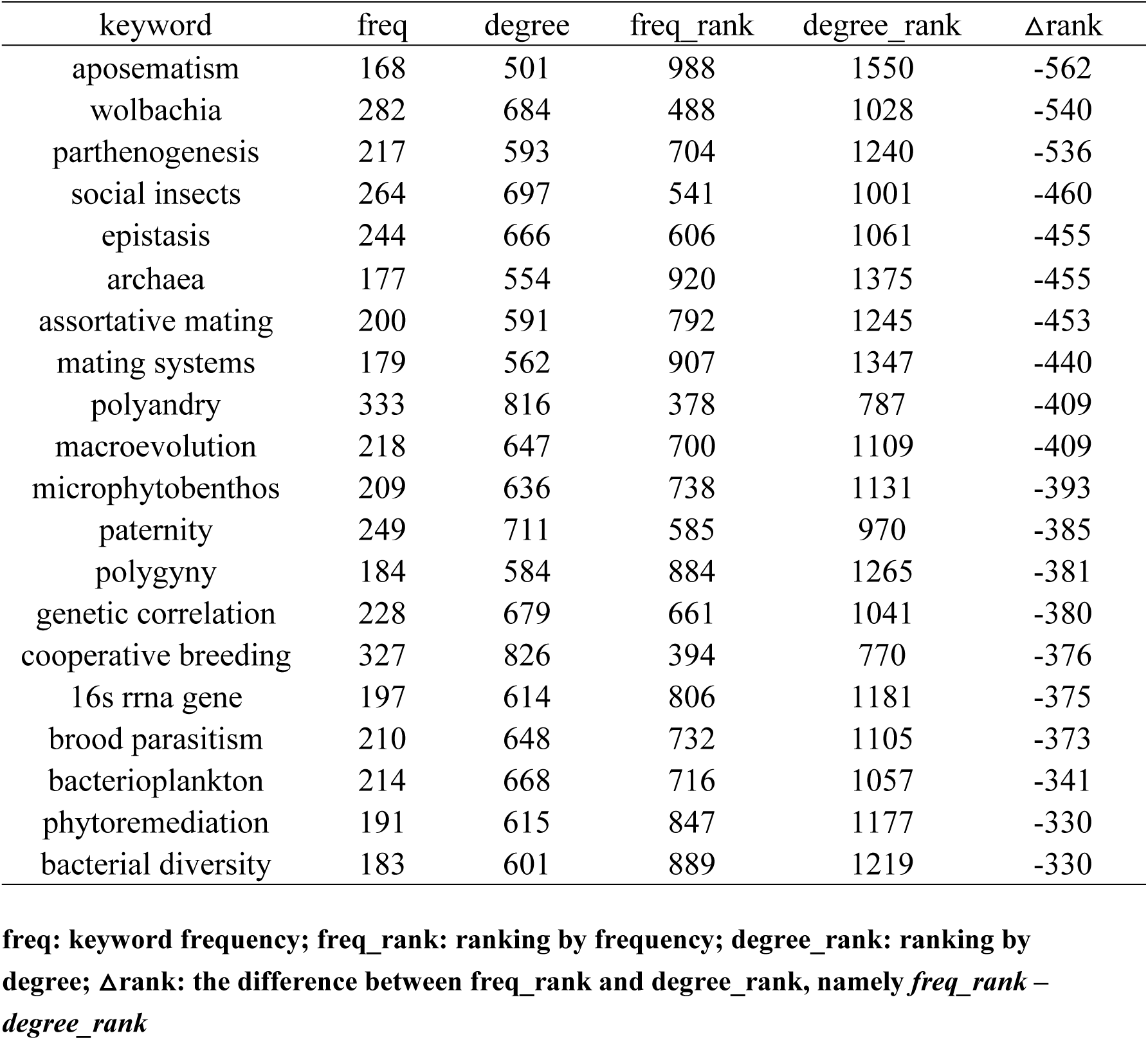
Top 20 keywords that tend to be overestimated by frequency

**Table 2.** Top 20 keywords that tend to be underestimated by frequency

| keyword | freq | degree | freq_rank | degree_rank | Δrank |
| --- | --- | --- | --- | --- | --- |
| semiochemicals | 123 | 713 | 1469 | 967 | 502 |
| plant population and community dynamics | 129 | 750 | 1390 | 897 | 493 |
| bayesian analysis | 145 | 779 | 1192 | 843 | 349 |
| monoterpenes | 143 | 759 | 1220 | 882 | 338 |
| gc-ms | 143 | 758 | 1220 | 884 | 336 |
| determinants of plant community diversity and structure | 170 | 922 | 972 | 643 | 329 |
| chemical ecology | 141 | 724 | 1249 | 945 | 304 |
| el niño | 140 | 716 | 1256 | 962 | 294 |
| conservation biogeography | 158 | 822 | 1065 | 781 | 284 |
| historical ecology | 143 | 731 | 1220 | 936 | 284 |
| invasion ecology | 163 | 835 | 1030 | 760 | 270 |
| long-term monitoring | 159 | 808 | 1056 | 793 | 263 |
| kairomone | 146 | 735 | 1184 | 930 | 254 |
| path analysis | 208 | 1094 | 746 | 496 | 250 |
| bioassay | 197 | 1018 | 806 | 558 | 248 |
| resource limitation | 145 | 719 | 1192 | 956 | 236 |
| autocorrelation | 157 | 779 | 1078 | 843 | 235 |
| bayesian | 193 | 965 | 831 | 607 | 224 |
| water availability | 147 | 722 | 1172 | 952 | 220 |
| field experiment | 250 | 1284 | 584 | 366 | 218 |

freq: keyword frequency; freq\_rank: ranking by frequency; degree\_rank: ranking by degree; $\Delta$ rank: the difference between freq\_rank and degree\_rank, namely $freq\_rank - degree\_rank$

### 2.5. Measuring keyword popularity in temporal dynamical network

In reality, ecological knowledge network was not built up in one step like we did in computer program, but growing brick by brick over time. Therefore, the knowledge network was not static, but temporal dynamical. Among the various network growing mechanisms, preferential attachment and node fitness might be two of the simplest ones, simple but useful. Preferential attachment, also known as “rich get richer” phenomenon, believes that pioneers with large degree have an advantage over newcomers and are more likely to form connections to other nodes in the future (Barabási & Albert 1999). On the other hand, node fitness, which is often described as “fit get richer” phenomenon, illustrates that newcomers could occasionally surpass the pioneers when they are intrinsically more attractive (Bianconi & Barabási 2001). We believed the combination of these two mechanisms could describe the dynamic patterns in our ecological knowledge network. Ecological topics being mentioned numerous times had solid theoretical basis or practical experience accumulation, thus are more likely to be included as keywords in the future. Nevertheless, new ecological topics never stop challenging the old ones and be ready to take their places in the field of ecological disciplines. This hypothesis led us to do the joint estimation of preferential attachment and node fitness in our ecological knowledge network, which would help us measure the keyword popularity more appropriately.

PAFit, a Bayesian statistical method, was used to estimate preferential attachment function and node fitness non-parametrically (Thong *et al.* 2016). In this method, the probability **P**_**i**_ for node **v**_**i**_ to get a new edge in the future is proportional to the product of attachment function **A**_**ki**_ and the fitness of the node **η**_**i**_: **P**_**i**_ ∝ **A**_**ki**_ × **η**_**i**_. The attachment function **A**_**k**_ = **k**^**α**^, where **k** is the degree of the node, and **α** is called attachment component. With the edge list with temporal information, the global attachment component **α** and fitness of each node **η**_**i**_ could be estimated non-parametrically. R package ‘PAFit’ was used to complete the whole task. Mathematical background and the application of the package could be found in Pham *et al.* 2017.

For our case, the product of attachment function and node fitness was calculated, this product (called as PAFit in our study) is used to measure the popularity of the keywords in the network. Due to the consideration of “rich-get-richer” and “fit-get-richer” phenomenon, PAFit is supposed to be superior to other simple metrics such as frequency and degree. However, this hypothesis should not be self-testifying but supported by facts. Therefore, we design the following experiment to verify our assumption.

### 2.6. Comparison of the predictive ability of frequency, degree and PAFit when measuring keyword popularity

To perform our experiment, we should answer a vital question in the first place: What is popularity? In the dictionary, popularity is “the quality or state of being popular” (“Popularity.” Merriam-Webster.com), while the definitions of popular include “of or relating to the general public” and “frequently encountered or widely accepted” (“Popular.” Merriam-Webster.com). Therefore, a popular keyword should be related to large amount of other keywords and occurring frequently in the ecological journals. These two characters could be well represented by degree and frequency mentioned in the former section.

Popularity of keywords should not only be descriptive but also predictive. In other words, when we say a keyword is popular, it has been popular for some time, and this trend will not disappear in the near future. For instance, if we gain the popularity of keywords in a specific time period, we might be able to predict the growth of the keywords in the following years. Therefore, we split our data into two parts, and tried to use the historical keyword popularity to predict the growth of keywords’ frequency and degree in the coming three years. The experiment procedure was designed as follows: 1. Construct the ecological knowledge network with data from 1988 to 2014, and calculate the frequency, degree and PAFit for every keyword appeared in these 27 years; 2. Construct the ecological knowledge network with data from 1988 to 2017, calculate the frequency and degree for every keyword appeared in the total 30 years; 3. Subtract the frequency of 27 years from frequency of 30 years, and we gain the change (or growth) of frequency in the recent three years (namely 2015-2017). The same is done to the keywords’ degree. Note that keywords emerging in the recent three years but not in the previous 27 years would be excluded from our analysis; 4. Fit a simple linear regression model using frequency, degree and PAFit in the former 27 years to predict the growth of frequency and degree in the following 3 years respectively. Compare the results and see if PAFit yields better predictions.

### 2.7. Commonality analysis to clarify relations of popularity metrics

This analysis was based on the regression models we got in the former section. Instead of using one metric at a time, we could include all three metrics and run a multiple regression. Obviously, the three metrics we compared are closely related to each other. Therefore, in the task of predicting the frequency growth and degree growth, they would share some explanatory power while each metric has its unique explanatory power. Commonality analysis is capable of decomposing the variance of R^2^ into unique and common variance of predictors. Though we did not intend to actually implement multiple regression to gain a better prediction of the popularity, this analysis could help us better understand the correlations among the three metrics. For instance, when we used PAFit to measure popularity, we got an adjusted R^2^, if adding frequency to do multiple regression was not going to rise up overall R^2^, then PAFit might contain enough power to depict popularity. In another way, when we have the R^2^ yielded by the frequency alone, and we found that including PAFit could promote the overall R^2^, then we could conclude that PAFit contains some explanatory power that frequency could not offer. Results of this analysis is showed in discussion. Detailed information about the method could be found in the previous study (Ray-Mukherjee *et al.* 2014). R packages ‘yhat’(Nimon *et al.* 2013) and ‘vegan’(Oksanen *et al.* 2013) were used to complete the tasks of calculation and visualization in commonality analysis.

## 3. Results

### 3.1. Overview of ecological knowledge network

From 1988 to 2017, the network density had decreased from 1.82×10^−3^ to 2.51×10^−4^(Fig.4A), which showed that the possibility for any two ecology-related keywords to co-occur in the same article was dropping in the recent three decades. Pearson correlation analysis showed that annual network density was negatively correlated with the distinct keyword number occurring in each year (r = −0.85, P < 0.01). The reason of the dropping density these years might be the exploding article number which brought numerous different keywords into the ecological area (Fig.1).

**Fig. 4.**
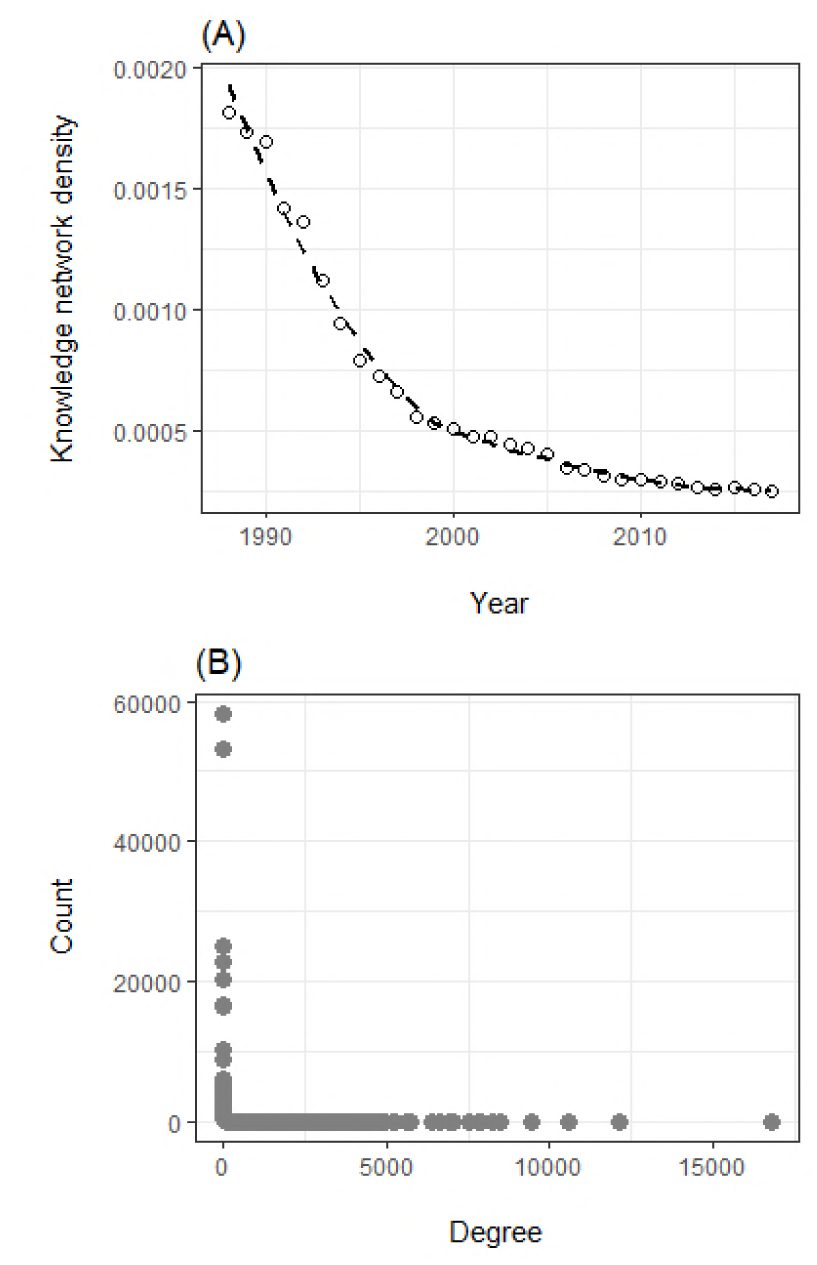
Basic property of the ecological knowledge network. (A) Temporal change of network density. (B) Degree distribution of the network.

Focusing on the degree distribution of the network, we found that it followed a power law distribution with a long tail, which indicated that very few nodes had extremely large amount of connections. It indicates that only few keywords could be enlisted time after time in the keyword area in ecological journals, while others appeared only once and never showed up again. Digging deeper, we could find that the point at the far right was the keyword “climate change”. With an occurrence number of 6,939, it was able to co-occur with 16,775 different keywords in the same article, and the penultimate point at the right is “biodiversity”, occurring 4,975 times and was related to 12,113 different keywords. On the other hand, it was found that 212,514 keywords had occurred only once and 38,018 occurred only twice. For these words, they could only co-occur with the keywords appearing in their same articles, therefore possessed a quite low degree (but not one, unless the article contained only one keyword). In such a background, if we could grasp the very few keywords with the highest degree, it’s possible for us to get a rather clear picture about the most popular topics in ecology.

### 3.2. Possible risks when using frequency to measure popularity in keyword analysis

Frequency had long been used to measure the popularity of topics in keyword analysis. Nevertheless, a keyword could have a large frequency simply for the reason that more papers about this topic were published in the investigated period, while other keywords might have relatively lower frequency but still be capable of making various links to different topics in the discipline. Inspecting the keywords with relatively higher frequency but lower degree, we could find that frequency tend to overestimate the popularity of ecological topics in microcosmic scale. In Table 1, the top 20 overestimated keywords were showed, we could find “aposematism” at the top of the list, which is a concept in evolutionary ecology, followed by “wolbachia” (all keywords were displayed in lower case), coming from subfield of microbial ecology. Take a further step, we found that the main sources of articles containing the top 20 keywords in this list were Evolution (462 articles containing at least one of these keywords), Proceedings of The Royal Society B: Biological Sciences (437), Behavioral Ecology and Sociobiology (363), Journal of Evolutionary Biology (353), FEMS Microbiology Ecology (336) and Molecular Ecology (332).

On the contrary, keywords related to macroscopic ecology tended to be underestimated by frequency metric, including words like “plant population and community dynamics”, “determinants of plant community diversity and structure”, “el niño”, “conservation biogeography” and “invasion ecology”(Table 2). Researches of macroscopic ecology are usually supported by large-scale spatial-temporal observations, which demands longer research cycle. This would definitely decrease the quantity of papers in the subfield, and consequently decrease number of relevant keywords. Interestingly, we found two other sorts of keywords that tend to be underestimated by frequency. One is keywords related to chemical ecology, including “semiochemicals”, “monoterpenes” and “kairomone”. It seemed that chemical ecology has a great potential to be applied in different aspects of ecology, while the paper volume in this subfield might be relatively low currently. The other was keywords related to methods in ecology and evolution, including “bayesian analysis”, “gc-ms” and “field experiment”. Among these words, “gc-ms” is closely related to chemical ecology, while “field experiment” is usually implemented on studies concerning macroscopic ecology. What we should notice is that as a challenger of frequentist statistics, Bayesian statistics has now gained its popularity in ecology. However, this popularity might be underestimated if we only focus how many times this keyword occurred in the previous literatures.

All in all, though frequency is always positively correlated with degree (in our case, we got a Pearson correlation coefficient of 0.98, P < 0.01), using it alone might misestimate the keyword popularity, and degree metric yielded based on the knowledge network could provide good supplementary information to fill the gap.

### 3.3. Measuring keyword popularity in a temporal dynamical network using PAFit

In Table 3, we could find that popularity metrics from the past 27 years could welly predict the growth of frequency and degree in the following 3 years (with R^2^ all larger than 0.75). The frequency metric performed better than degree at predicting the future growth of frequency (R^2^ = 0.82 > 0.77), while the degree metric surpassed frequency at predicting the future growth of degree (R^2^ = 0.79 > 0.76). However, both metrics were beat by PAFit, no matter in frequency growth prediction or degree growth prediction (R^2^ reached 0.89 in both tests).

**Table 3.** Comparison of performance when using simple linear regression to predict the keyword popularity by different metrics

| Predictor | Predicting Δfrequency | | Predicting Δdegree | |
| --- | --- | --- | --- | --- |
| | Formula | $R^2$ | Formula | $R^2$ |
| Frequency | $y = -0.20 + 0.25x$ | 0.82 | $y = 0.80 + 0.66x$ | 0.76 |
| Degree | $y = -0.59 + 0.07x$ | 0.77 | $y = -0.42 + 0.20x$ | 0.79 |
| PAFit | $y = -0.53 + 0.11x$ | 0.89 | $y = -0.21 + 0.30x$ | 0.89 |

Ranking the keywords from the total 30 years’ data according to PAFit, we could detect the ecological hotspots in the recent three decades (Table 4). The top 10 ecological topics in descending order were “climate change”, “biodiversity”, “invasive species”, “conservation”, “ecosystem services”, “dispersal”, “species richness”, “competition”, “functional traits” and “disturbance”. It was noteworthy that “invasive species”, “ecosystem services” and “functional traits” have relatively lower frequency and degree among the top 10 keywords, however, their intrinsic fitness (η) were very high, which indicates that there are great chances for these topics to become more prevalent in the future.

**Table 4.**
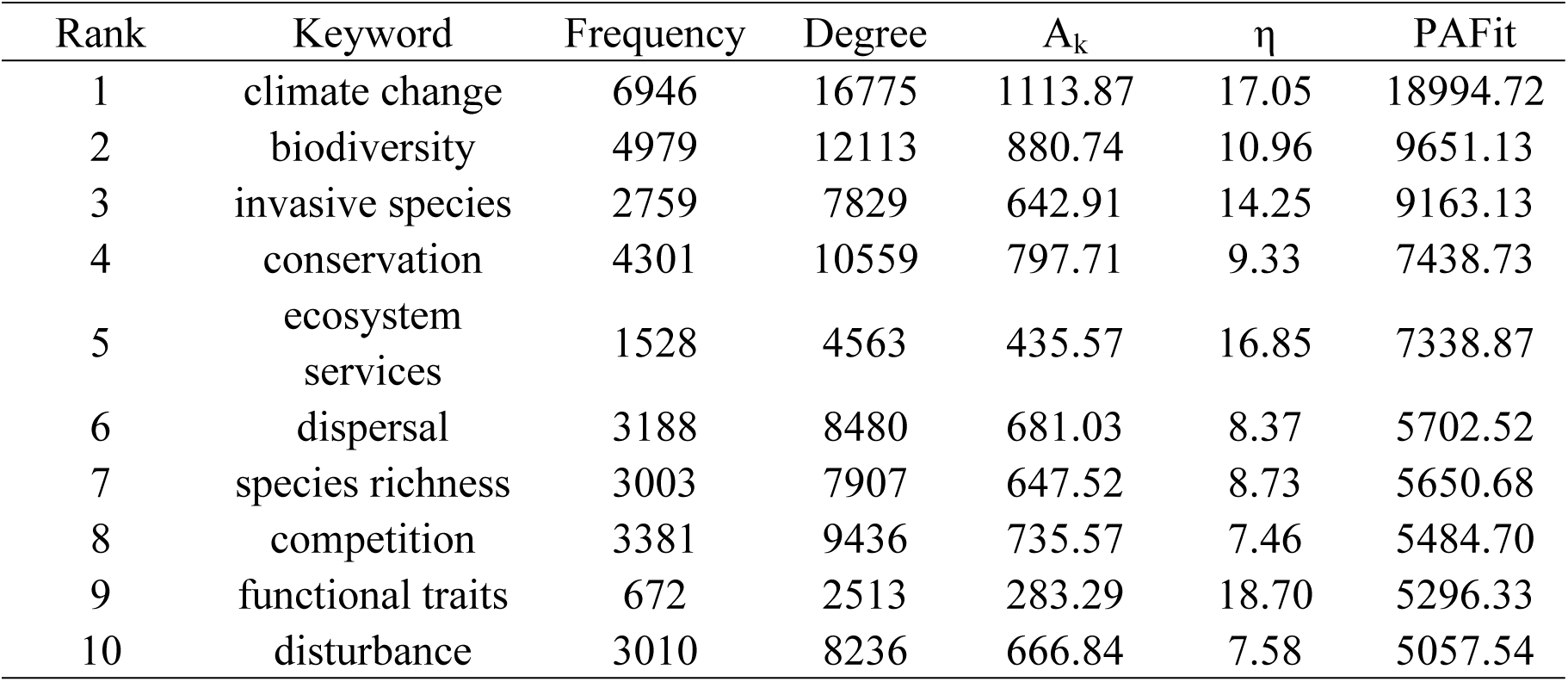
Top 10 ecological hotspots ranked by PAFit

## 4. Discussions

### 4.1. Strong correlations between metrics discussed in our study

In our study, we have used three metrics to measure the popularity of ecological topics, namely frequency, degree and PAFit. In essence, the growth of degree is a sufficient but not necessary condition for the growth of frequency. That is to say, when the degree of a keyword rises, the frequency would definitely increases. Nevertheless, the opposite might not be true when the keyword is related to merely several keywords in its subfield. According to our results, some topics in microcosmic ecology could gain a relatively high frequency due to the average short research cycle. That is why degree could be a good supplementary metric to frequency. And when we consider the popularity of keywords in a network, we noticed that the “rich-get-richer” and “fit-get-richer” phenomenon did exist in our temporal network. This was testified by the superior performance of PAFit in predicting the growth of frequency and degree, beating the frequency and degree metrics themselves. But take a step backward and we could find that the three metrics discussed in our study are obviously correlated with each other. For one, frequency of a keyword could also be interpreted as how many articles containing a specific ecological topic were published in the investigated period. The more the frequency, the more likely that this ecological topic could be related to other ecological topics. Therefore, there is a statistically strong positive correlation between frequency and degree in most cases. On the other hand, when consider things in a network, degree is actually a component of PAFit. As the equation of PAFit could be displayed as: PAFit = k^α^ × η, where k is the degree, α is the attachment component and η is the node fitness. When we make α=1, η=1, this becomes equivalent to degree. Technically speaking, using degree to measure popularity is a specific case of PAFit, where we make assumptions that node fitness mechanism does not exist and the attachment component equals to 1. This model had been discussed and the pattern was coined as “scale-free feature” in 1999 by Barabási and Albert, and PAFit was a developed model built on this. So should we use PAFit alone to measure keyword popularity? The technical answer might be yes. If we define popularity the same way as mentioned in our method, then we could do a commonality analysis to clarify the relations among the three metrics. When predicting the frequency growth, if we already include PAFit in the model, adding degree and frequency could only promote 3.36% of the total adjusted R^2^ (Table 5), and this promotion reduced to 0.40% when predicting the degree growth (Table 6). The overlapping area of variance commonly explained by the three metrics reached 0.79 and 0.76 for predicting frequency growth and degree growth respectively (Fig.5). This is already a great amount, which means that frequency alone could grasp the most general trends in keyword analysis. However, the explained variance brought by PAFit (0.10 predicting frequency growth and 0.11 predicting degree growth) was irreplaceable and could make a real difference in the popularity measurement.

**Fig. 5.**
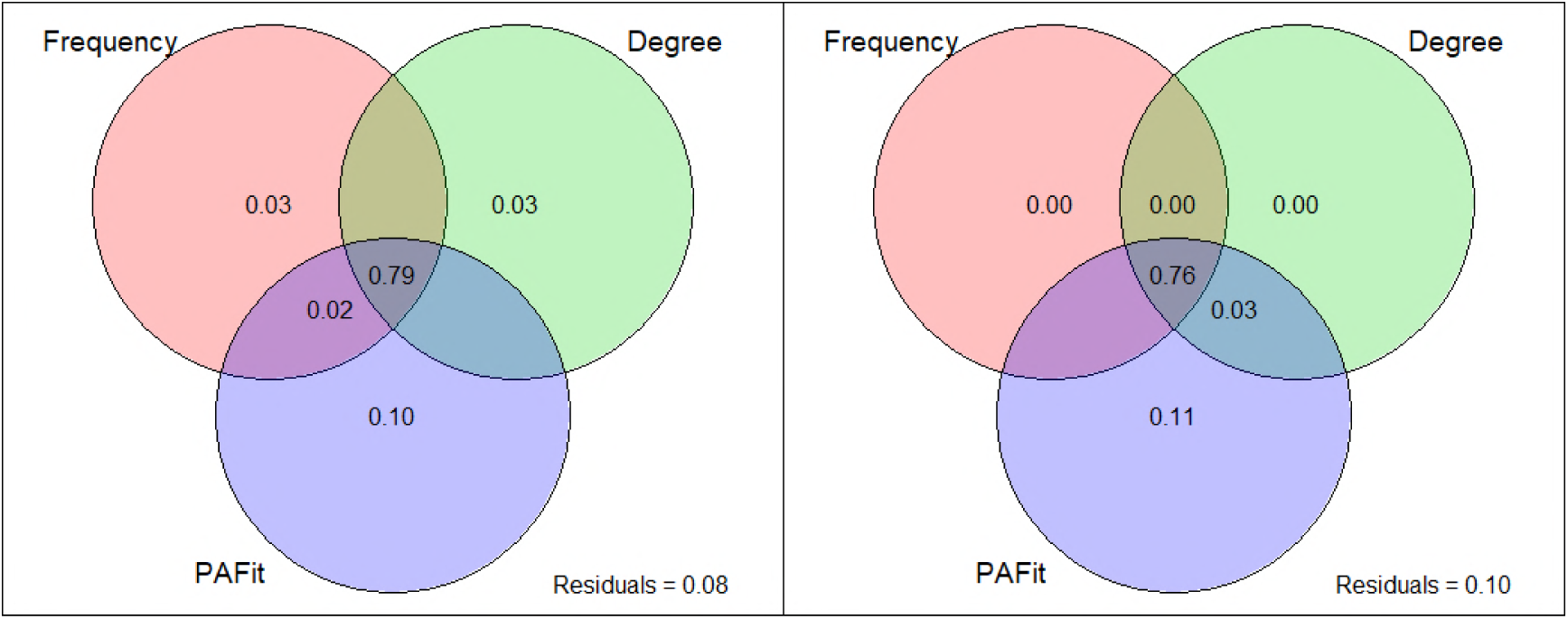
Using variation partition analysis to clarify the explainable variance among three metrics (frequency, degree and PAFit) when predicting the changes of frequency (left) and degree (right).

**Table 5.** Partition table of variance when predicting the change of frequency

|  | Ajusted R<sup>2</sup> | %Total |
| --- | --- | --- |
| Frequency | 0.816 | 88.55% |
| Degree | 0.768 | 83.38% |
| PAFit | 0.890 | 96.64% |
| Degree + Frequency | 0.817 | 88.73% |
| Frequency + PAFit | 0.892 | 96.81% |
| Degree + PAFit | 0.895 | 97.17% |
| Degree + Frequency + PAFit | 0.921 | 100.00% |

**Table 6.** Partition table of variance when predicting the change of degree

|  | Ajusted R<sup>2</sup> | %Total |
| --- | --- | --- |
| Frequency | 0.765 | 84.95% |
| Degree | 0.793 | 88.04% |
| PAFit | 0.897 | 99.60% |
| Degree + Frequency | 0.793 | 88.06% |
| Frequency + PAFit | 0.901 | 99.96% |
| Degree + PAFit | 0.899 | 99.74% |
| Degree + Frequency + PAFit | 0.901 | 100.00% |

Nevertheless, in practice frequency and degree are more intuitional indexes than PAFit. Frequency is the number of articles containing the keyword, degree is the number of keywords that co-occur with the keyword in the same article. PAFit is a metric that could be used to measure the probability of the keyword to co-occur with other keywords, which could be a little abstract to understand. Therefore, we believe that PAFit is the best metric to use when we try to measure keyword popularity, but frequency and degree should always be provided as supplementary metrics so that we could explain our results more intuitively.

### 4.2. The latent capability of node fitness to detect potential ecological hot topics

Previous discussion had shown that PAFit could totally replace frequency and degree when our task was to predict keywords’ popularity, and the unique variance that it surpasses the other two metrics actually comes from the special consideration of node fitness. Node fitness could explain why late-comers could surpass first-movers, which would never happen in rich-get-richer mechanism. Previous study had used node fitness to measure the competitiveness of authors in a citation network (Ronda-Pupo and Pham 2018). It was observed that some late-comers acquired even more citations than the first-movers in scientific publication (Newman 2009). The main reason was interpreted as the fitness could reflect the qualities of the authors’ scientific contributions. In our case, the keyword fitness reflects the innate popularity of an ecological topic. Some ecological topics did not appear until very late in the disciplinary history, while others might be coined but not prevailed then. But when these topics meet the needs of time, they could get hot in a rather short period. For instance, the concept of “ecosystem services” had been suggested in late 2000s, but it did not gain a real leap in popularity until the monumental work Millennium Ecosystem Assessment was published in 2005(Fisher et al. 2009).

According to our study, we could find that node fitness had weak correlations with other metrics (Table 7), which indicates that it has a potential to offer new explainable power for the invisible popularity of ecological topics that usually neglected by the common view. We had used frequency growth and degree growth to reflect the keyword popularity, but when we take growth rate (divide growth by the original number of frequency or degree) into consideration, we found that fitness is more correlated to frequency growth rate and degree growth rate than other metrics. Based on our research data, we made a list of the top 10 potential ecological hotspots based on node fitness(Table 8). Compared with the hotspots we found using PAFit (Table 4), we could find that some of fittest keywords had already gained much popularity, including “functional traits”, “climate change” and “ecosystem services”. Moreover, it seems that molecular technology has great potential to develop the discipline of ecology, with many potential hot topics like “metabarcoding”, “high-throughput sequencing”, “next-generation sequencing”.

**Table 7.**
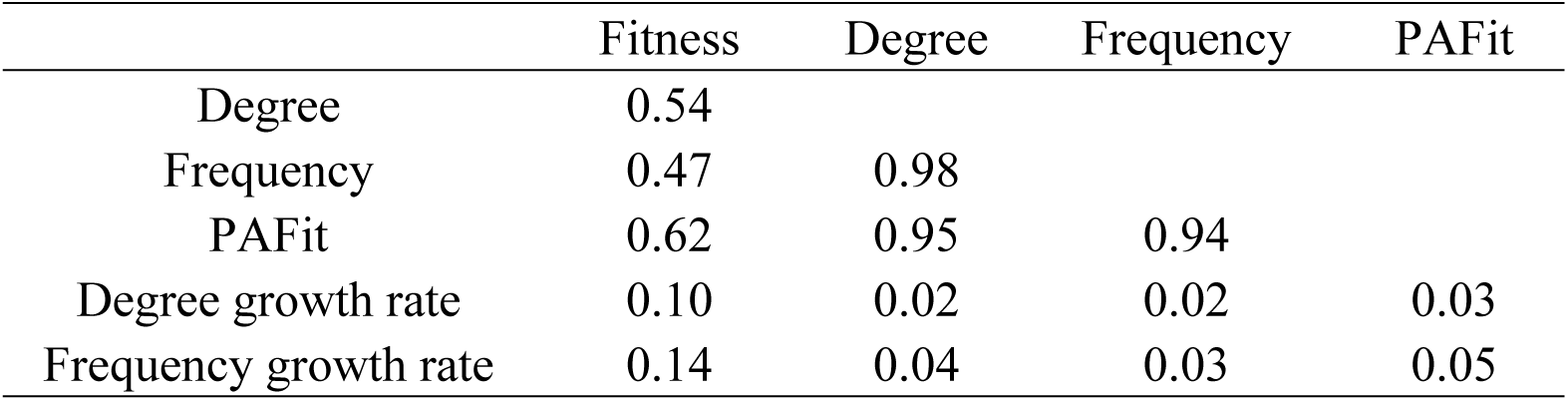
Correlations among popularity metrics and their correlation with degree and frequency growth rate

|  | Fitness | Degree | Frequency | PAFit |
| --- | --- | --- | --- | --- |
| Degree | 0.54 |  |  |  |
| Frequency | 0.47 | 0.98 |  |  |
| PAFit | 0.62 | 0.95 | 0.94 |  |
| Degree growth rate | 0.10 | 0.02 | 0.02 | 0.03 |
| Frequency growth rate | 0.14 | 0.04 | 0.03 | 0.05 |

**Table 8.** Top 10 ecological hotspots ranked by keyword fitness

| rank | word | fitness |
| --- | --- | --- |
| 1 | functional traits | 18.70 |
| 2 | climate change | 17.05 |
| 3 | ecosystem services | 16.85 |
| 4 | metabarcoding | 16.36 |
| 5 | citizen science | 16.23 |
| 6 | high-throughput sequencing | 16.10 |
| 7 | environmental filtering | 15.85 |
| 8 | next-generation sequencing | 15.43 |
| 9 | species distribution model | 15.41 |
| 10 | cultural ecosystem services | 15.07 |

### 4.3. Application of egocentric network analysis to explore the trends in subfields

In bibliometric study, keyword analysis are commonly used to analyze the trend of a specific research area, and frequency are often used as the only criteria to quantify keyword popularity (Aleixandre-Benavent et al. 2018, Romanelli et al. 2018, Yin et al. 2018). After the calculation of frequency, keywords are ranked and the top keywords are selected to reflect the research hotspots. Our study showed that PAFit is a better metric to measure keyword popularity, because it has considered both accumulative advantage and innate attractiveness of topics represented by keywords. However, another important point should not be neglected, that is we considered ecological topics were related in a knowledge network. In our study we had tested our assumptions using all the information we had in the selected ecological journals. But if we were only interested in a subfield in ecology, we could easily extract the relevant data and establish a local network, so as to explore the trends in the subfield.

In social science, egocentric network analysis has been widely used to understand individuals and their immediate social environment (Wu et al. 2016, Perry et al. 2018). Ego network consists of a focal node (“ego”) and nodes that directly connected to it (“alters”). When it comes to our ecological knowledge network, constructing ego networks could help us dig deep into a subfield. For example, if a research team focuses on doing ecological research using remote sensing, they might take interests in the existing hotspots and potential hot topics. In this way, we could build an ego network with the focal keyword “remote sensing” (Fig.6). All the keywords appearing in the network had been co-occurred with “remote sensing” in the same article at least once. In the local scale, “remote sensing” tend to co-occur more with keywords “climate change”, “biodiversity”, “conservation”, “species richness” and “disturbance” (displayed in triangular nodes). In the global scale, “climate change”, “conservation”, “ecosystem services”, “biodiversity” and “invasive species” were the most popular among topics related to remote sensing in ecology (nodes in red), and the top 5 potential hot topics were “climate change”, “ecosystem services”, “plant-plant interactions”, “functional traits” and “citizen science”. Topics like “climate change” had been popular already and are going to be even more popular in the future, researchers in this subfield had recognized its importance and lots of studies had performed on this topic. Topics like “citizen science”, on the other hand, were rarely mentioned in ecology and there were relatively fewer researches concerning both remote sensing and citizen science at the moment, but there’s great hope that citizen science would be combined with remote sensing and make great contributions to the development of ecology in the future.

**Fig. 6.**
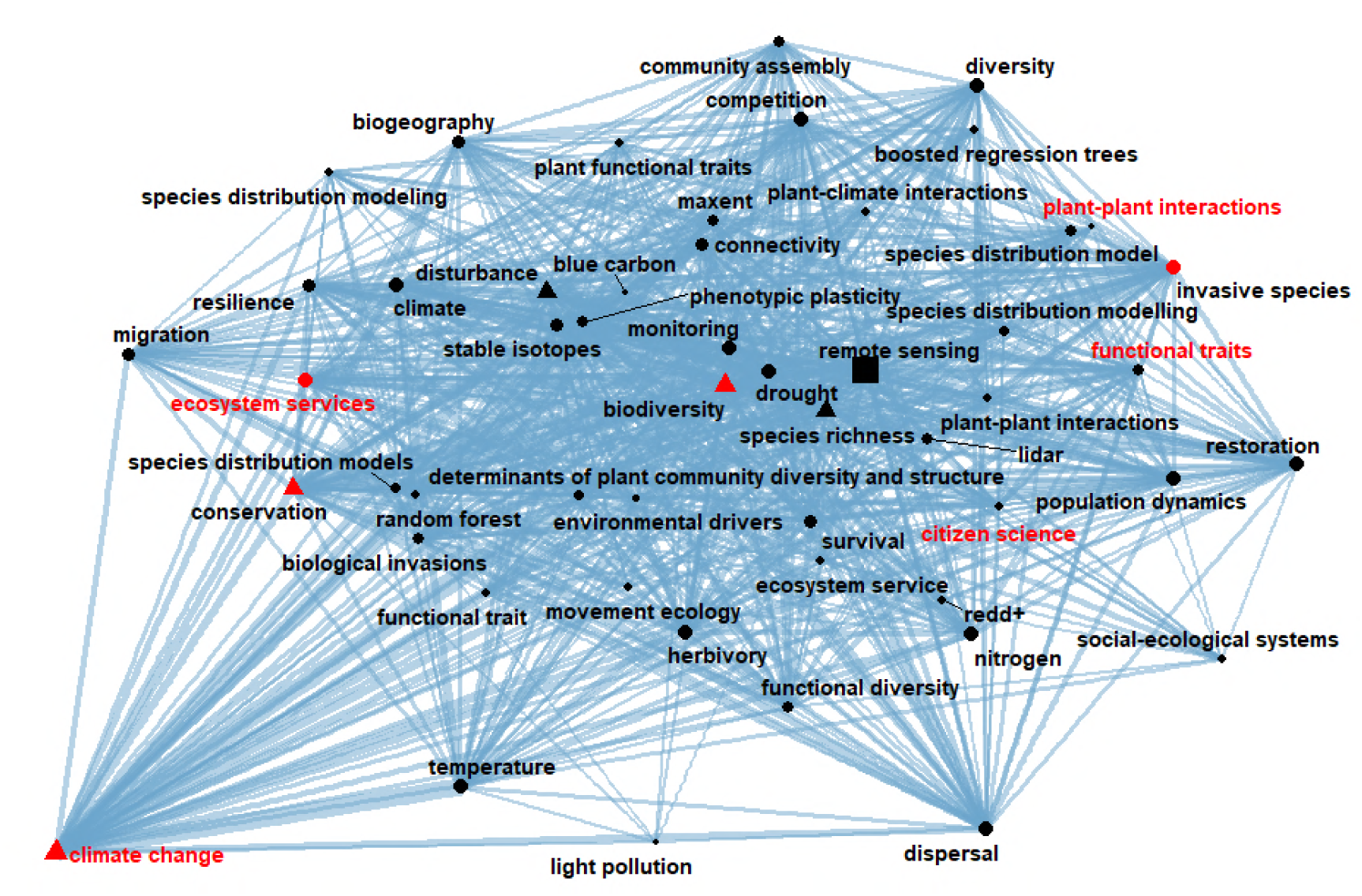
Egocentric network analysis for “remote sensing”. The square node in the middle is “remote sensing”. Sizes of nodes are proportional to the local degree of the nodes in the ego network, and the top 5 local popular keyword are in the shape of triangle. Width of edges are proportional to the number of co-occurrence between keywords. Nodes are selected according to their PAFit and fitness in the complete network, top 30 fittest and top 30 most popular keywords are chosen to establish the network. Nodes in red are top 5 popular keywords, nodes with red labels are top 5 fittest keywords.

## 5. Conclusions

In our study, we have displayed our ecological knowledge structure in the form of network, which enables us to better quantify the popularity of ecological topics. This will definitely promote our comprehension on the whole discipline as well as development in every subfield of ecology. Ecological knowledge network could be constructed to depict the ecological development in different time ranges, different regions and different domains, and considering the abundant achievements in graph theory and various applications in network analysis, more interesting discoveries could be found in ecological knowledge network. In the era of “big literature”, with large amount of accessible data and all sorts of digital tools at hand, we are capable of drawing a tremendous map of our ecological world. We believe this map could give us a clearer picture of our discipline, and guide us to more collaborations, deeper discipline integration and better researches in the future.

